# Glutamine is essential for overcoming the immunosuppressive microenvironment in malignant salivary gland tumors

**DOI:** 10.1101/2022.04.29.490103

**Authors:** Shuting Cao, Yu-Wen Hung, Yi-Chang Wang, Yiyin Chung, Yue Qi, Ching Ouyang, Xiancai Zhong, Weidong Hu, Alaysia Coblentz, Ellie Maghami, Zuoming Sun, H. Helen Lin, David K. Ann

**Author notes:** To whom all correspondence should be addressed: David K. Ann, Ph.D., Beckman Research Institute, City of Hope Comprehensive Cancer Center, Duarte, CA 91010-3000. Contribute equally (Yu-Wen Hung, Yi-Chang Wang, Yiyin Chung, Yue Qi).

## Abstract

**Rationale:** Immunosuppression in the tumor microenvironment (TME) is key to the pathogenesis of solid tumors. Tumor cell-intrinsic autophagy is critical for sustaining both tumor cell metabolism and survival. However, the role of autophagy in the host immune system that allows cancer cells to escape immune destruction remains poorly understood. Here, we determined if attenuated host autophagy is sufficient to induce tumor rejection through reinforced adaptive immunity. Furthermore, we determined whether dietary glutamine supplementation, mimicking attenuated host autophagy, is capable of promoting antitumor immunity.

**Methods:** A syngeneic orthotopic tumor model in *Atg5*^*+/+*^ and *Atg5*^*flox/flox*^ mice was established to determine the impact of host autophagy on the antitumor effects against mouse malignant salivary gland tumors (MSTs). Multiple cohorts of immunocompetent mice were used for oncoimmunology studies, including inflammatory cytokine levels, macrophage, CD4^+^, and CD8^+^ cells tumor infiltration at 14 days and 28 days after MST inoculation. *In vitro* differentiation and *in vivo* dietary glutamine supplementation were used to assess the effects of glutamine on Treg differentiation and tumor expansion.

**Results:** We showed that mice deficient in the essential autophagy gene, *Atg5*, rejected orthotopic allografts of isogenic MST cells. An enhanced antitumor immune response evidenced by reduction of both M1 and M2 macrophages, increased infiltration of CD8^+^ T cells, elevated IFN-γ production, as well as decreased inhibitory Tregs within TME and spleens of tumor-bearing *Atg5*^*flox/flox*^ mice. Mechanistically, ATG5 deficiency increased glutamine level in tumors. We further demonstrated that dietary glutamine supplementation partially increased glutamine levels and restored potent antitumor responses in *Atg5*^*+/+*^ mice.

**Conclusions:** Dietary glutamine supplementation exposes a previously undefined difference in plasticity between cancer cells, cytotoxic CD8^+^ T cells and Tregs.

## Introduction

Accumulating evidence suggests that the modulation of immune microenvironment plays a critical role in anti-cancer immunity by regulating both tumor immune surveillance and evasion [reviewed in [1-3]]. Notably, the immunosuppressive networks promote cancer progression, metastasis and resistance to therapies [1]. Salivary gland tumors have more than 30 subtypes, among them, salivary duct carcinoma (SDC), albeit rare, represents the most lethal and aggressive histologic subtype [4]. A recent study by Linxweiler et al. compared the immune landscape of malignant salivary gland tumors (MSTs) and revealed a significantly higher overall immune score [5]. In addition, several groups reported that MSTs exhibit higher levels of cytotoxic T lymphocyte (CTL) dysfunction (epitomized by overexpression of immune checkpoint genes) and an abundance of immune-suppressive cell types (exemplified by regulatory T cells [Tregs]) [6-10]. Thus, MSTs have evolved strategies to module the immune microenvironment to evade antitumor immune response.

Accumulating evidence has suggested the involvement of the various nutrients in regulating the survival, apoptosis, differentiation, activation, effector function and tumor trafficking of immune cell subsets [11-13]. For example, during T-cell differentiation, CD4^+^ and CD8^+^ cells are generated from naïve T cells through distinct glucose-mediated activation of effector function and clonal selection [12]. CD8^+^ T cells are the preferred immune cells for targeting cancer cells for immunogenic cell death [14, 15]. In parallel, the naïve T cells, activated by antigen-presenting cells and specific cytokines, differentiate into CD4^+^ effector cells (T helper cells and Th17 cells) as well as Tregs [11, 16]. Notably, Treg cells exhibited increased fatty acid oxidation, whereas Th17 cells have demonstrated a reliance upon fatty acid synthesis [17]. Tregs appear to play a major role in suppressing antitumor immune responses [18]. The precise nutrient utilization pathways regulating Treg functions and the crosstalk between different T lymphocyte subsets to govern antitumor immunity remains unclear.

Autophagy recycles cargos to provide anabolic and catabolic substrates [19]. This metabolic recycling function of autophagy promotes tumor cell survival under conditions of nutrient limitation [20]. Furthermore, autophagy may favor tumor progression by promoting the escape of malignant cells from immune surveillance [21-25]. Autophagosome formation during autophagy involves various autophagy-related genes (Atgs), including *Atg5* [26]. Indeed, elevated *Atg5* expression is an unfavorable prognostic marker for human renal and hepatic cancers (The Human Protein Atlas; [27]). Moreover, autophagy plays a key role in shaping T cell immunity and activation [28, 29]. During the process of activation and differentiation to effectors, T cells undergo metabolic reprogramming and shift from anabolic to catabolic mode [30].

Autophagy has emerged as a crucial regulator of T cell catabolic activity [31]. Deletion of *Atg7, Atg5* or *Atg3* impairs CD4^+^ and CD8^+^ T cell proliferation and function in knockout mice [28, 32, 33], whereas deletion of *Atg7* or *Atg5* leads to Treg depletion and greater antitumor response [34]. Autophagy also promotes invariant natural killer T (iNKT) cells and Tregs differentiation in the thymus [35]. Hence, autophagy regulates the dynamic nature of antitumor immunity and homeostasis.

To determine whether autophagy changes impact tumor progression, most reports were centered on how tumors exploit their intrinsic autophagy competency to survive antitumor immunity in the hostile tumor microenvironment (TME) [36, 37]. In contrast, our limited understanding of the effect of host autophagy on the function and integrity of immune mediators that promote tumor progression versus mediators that promote tumor rejection is mainly derived from *in vitro* immune cell culture systems and therefore limited. In other words, mechanisms by which host autophagy stimulates or limits the immune system for recognizing and fighting tumor cells in tumor rejection remain unclear [38, 39]. Here we utilized an *in vivo* model in which both autophagy-attenuated *Atg5*^*flox/flox*^ and autophagy-competent *Atg5*^*+/+*^ mice were orthotopically allografted with syngeneic MST cells [40] to examine the role of attenuated host autophagy in regulating antitumor immune response within TME. For the first time, we present evidence that autophagy was associated with a reduction in intratumor glutamine level and suppressed cytotoxic T lymphocyte activity, favoring MST progression. Lastly, dietary glutamine supplementation retarded tumor growth and enhanced host antitumor immunity. Our findings provide a rationale for dietary glutamine supplementation as a therapeutic strategy to exploit the metabolic vulnerability of T cells against MST.

## Material and Methods

### Mice breeding

All animal protocols were in accordance with the guideline of Institutional Animal Care and Use Committee at City of Hope (IACUC 06038). <u>Mice</u> were housed in a <u>specific pathogen-free</u> room with a 12-h light/dark cycles and were fed an autoclaved <u>chow diet</u> and water *ad libitum. LGL-KRAS*^*G12V*^ mice, Ela-CreERT mice, and *Atg5*^*flox/flox*^ mice [13] were crossed to derive *Ela-CreERT;LGL-KRAS*^*G12V*^*;Atg5*^*flox/flox*^ (*KRAS*^*G12V*^*;Atg5*^*flox/flox*^), and *Ela-CreERT;LGL-KRAS*^*G12V*^*;Atg5*^*+/+*^ (*KRAS*^*G12V*^; *Atg5*^*+/+*^) mice, as we described previously [40]. Genotyping was conducted as described previously [41, 42]. Adult male mice, 8-10 weeks of age, were used in all experiments.

### Diet

All mice were kept on normal chow until start of the experiments. Diets used in this study are based on the open standard diet with 16 %kcal fat with crystalline amino acids from Research Diets Inc. (New Brunswick, NJ, USA). The Control diet (A11112201) contained all essential amino acids and nonessential amino acids as specified by Research Diets. Glutamine-supplemented diet contained all amino acids equal to the control diet with the addition of 200 g of glutamine by Ishak Gabra [43]. Corn starch content was adjusted to achieve the isocaloric intake. Mice were fed with respective diets for 28 days. Glutamine concentration was determined in the collected serum and harvested submandibular glands (SMGs), respectively.

### Tumor digests and submandibular glands tumor cells isolation

Tumors were cut into small pieces and digested into single cell suspension as previously described [40]. The tumors were minced and digested up to 60 min at 37 ° in digestion medium containing collagenase (1 mg/ml; MilliporeSigma, C6885), hyaluronidase (100 units/ml; MilliporeSigma, H3506), DNase I (50 µg/ml; MilliporeSigma, D4527), bovine serum albumin (1 mg/ml; MilliporeSigma, A2153), HEPES (pH 7.3, 20 mM; Corning, 25-060-CI) in Dulbecco’s Modified Eagle’s Medium/Ham’s F-12 50/50 Mix (Corning, 16-405-CV). The suspension of digested tumor cells was passed through a 100 µm sieve to remove the remaining tissue chunks. The red blood cells were lysed by incubating cell suspensions in 1X red blood cell lysis buffer (155 mM NH_4_Cl, 12 mM NaHCO_3_, 0.1 mM EDTA, pH 7.3) for 3 min on ice.

### Primary tumor cell culture

The primary cells were plated on collagen I-coated dishes (Corning, 354450), and maintained in a medium consisting of Dulbecco’s Modified Eagle’s Medium (Corning, 10-013-CV) with fetal bovine serum (10%; Thermo Fisher Scientific, 10437028), L-glutamine (5 mM; Thermo Fisher Scientific, A2916801), hydrocortisone (400 ng/ml; MilliporeSigma, H0888), insulin (5 µg/ml; Thermo Fisher Scientific, 12585014), EGF (20 ng/ml; Thermo Fisher Scientific, PHG0311), HEPES (15 mM; Thermo Fisher Scientific, 15630080) and antibiotic-antimycotic (1X; Thermo Fisher Scientific, 15240112). After 1 to 2 week of incubation, colonies of the GFP-negative tumor cells were manually picked and transfer to new cell culture dishes.

### Orthotopic tumor implantation

To distinguish host genotypes from genotypes of inoculated tumor cells, the tumor cells collected from *KRAS*^*G12V*^; *Atg5*^*+/+*^ and *KRAS*^*G12V*^; *Atg5*^*flox/flox*^ tumor-bearing mice were designated as *KRAS*^*G12V*^*;Atg5*^*+/*+^ and *KRAS*^*G12V*^*;Atg5*^*Δ/Δ*^, respectively. Whereas host genotypes were designated as *Atg5*^*+/+*^ and *Atg5*^*flox/flox*^. *Atg5*^*+/+*^ and *Atg5*^*flox/flox*^ mice were orthotopically inoculated with 2×10^5^ MST cells (*KRAS*^*G12V*^*;Atg5*^*+/+*^ and *KRAS*^*G12V*^*;Atg5*^*Δ/Δ*^), suspended in DMEM/Matrigel (1:1), in the right (SMGs). Tumor sizes were measured at least three times a week with digital calipers and tumor volume was calculated using the formula Volume (mm^3^) = (W^2^ x L)/2, where W (width) and L (length) correspond to the smaller and larger of two perpendicular axes, respectively. Animals were euthanized post-tumor implantation at either early end-point (Day 14) or late end-point (Day 25) according to humane endpoints as specified by COH IACUC guideline. For high glutamine diet feeding experiments, 5×10^4^ MST cells were implanted in SMG of mice. Tumor-bearing mice were fed with regular or high glutamine diet for another 21 days. Diets were changed weekly, and the consumption of diets were measured. Tumor-bearing mice were euthanized at 21 days post-tumor implantation.

### LPS treatment

Naïve mice were intraperitoneal injected with 5 mg/kg body weight lipopolysaccharides (LPS; MilliporeSigma, LPS25) in PBS or equal volume of PBS. Six hours following LPS administration, spleens were harvested, and spleen weights measured.

### Tissue preparation and characterization

Tumors were excised and fixed in 10% neutral-buffered formalin (MilliporeSigma, HT501128) for 48 h. Tissue embedding, sectioning, and staining with modified Mayer’s hematoxylin (American MasterTech, HXMMHGAL) and eosin Y stain (American MasterTech, STE0157), or H&E stain, were performed in City of Hope Pathology Core as previous described [40].

### Immunohistochemistry and quantification

The immunohistochemistry (IHC) was performed by City of Hope Pathology Core as described previously [40-42]. Briefly, formalin fixed paraffin embedded (FFPE) tumor tissue slides were deparaffinized and hydrated through xylenes and graded alcohol solutions. The tissue slides were pressure-cooked in citrate-based unmasking solution for 30 min and washed in phosphate-buffered saline for 5 min, followed by quenching of endogenous peroxidase activity in H_2_O_2_ (0.3%; MilliporeSigma, H1009) for 30 min. The slides were then blocked for 20 min with a mixture of Avidin D solution and diluted normal blocking serum, which was prepared from the species in which the secondary antibody is made. The slides were then incubated with a mixture of primary antibody and biotin solution for 30 min and washed in buffer 3 times. The slides were incubated in the Vector biotinylated secondary antibody for 30 min, washed for 5 min, and then incubated in Vectastain Elite ABC Reagent for 30 min. After being washed for 5 min, the slides were processed with the DAB Substrate Kit. Primary antibodies for IHC include antibody recognizing Ki67 (abcam, ab15580), F4/80 (Bio-Rad, MCA497R), CD11b (Abcam, ab133357), CD4 (Biolegend, 201501), CD8 (Thermo Fisher, 14-0808-82). For quantification, 10x magnification images of 5 nonoverlapping fields of tumors (5 images per mouse) were quantified using Image-Pro Premier 9.02 (Media Cybernetics).

### qRT-PCR

RNA was extracted using Trizol (Invitrogen, 15596026) according to the manufacturer’s instructions. The concentration of the isolated RNA and the ratio of absorbance at 260 nm to 280 nm (A260/A280 ratio) were measured with spectrophotometer (Biotek). cDNA was generated using iScript Kit (Bio-Rad, 1708890) and the qRT-PCR reaction utilized the components contained in the iTaq Universal SYBR Green Supermix (Bio-Rad, 1725120). Household gene transcript levels (*Gapdh*) were used for normalization. The 2−ΔΔCt method was used to analyze the relative changes in each target gene expression [44]. Sequences of the primers are listed in Table S1.

### Immunoblotting

Whole tissue protein was extracted by Qproteome Mammalian Protein Prep Kit (Qiagen, 37901) according to the manufacture’s guidelines. Cell lysates were prepared by directly lysing cells in 1X Laemmli SDS-PAGE buffer with vortexing and heating at 95 °C for 10 min. The protein concentration was measured by Bicinchoninic acid assay. Western blotting was performed by running equal amount of protein on a SDS-PAGE gel and immunoblotted with primary antibodies of interest followed by horseradish peroxidase conjugated secondary antibody following manufacturer’s instruction. After chemiluminescent reaction, blots were visualized with a Chemi-Doc Touch Imaging System (Bio-Rad).

### Multicolor flow cytometry

Cell suspensions (splenocytes and SMG tumor cells) were stained in FACS buffer (PBS supplemented with 1% BSA) for 30 min on ice using the following antibodies: CD11b-APC-eFluor 780 (Invitrogen, 47-0112-80), F4/80-eFluor 450 (Invitrogen, 48-4801-80), MHCII-PerCP-eFluor 710 (Invitrogen, 46-5320-80), CD86-PE-Cyanine7 (Invitrogen, A15412), CD206-PE (Invitrogen, 12-2061-82), Ly6b-APC (Novus, NBP2-13077APC), NK1.1-Super Bright 702 (Invitrogen, 67-5941-82), CD49b-APC (Invitrogen, 17-5971-81), CD4-PE-Cyanine7 (Invitrogen, 25-0041-81), and CD8-eFluor 450 (Invitrogen, 48-0081-80), diluted in FACS buffer at 1:100 ratio, whereas CD25-SB600 (Invitrogen, 63-0251-80) diluted in FACS buffer at 1:50 ratio. For intracellular cytokine staining, cells isolated from *Atg5*^*+/+*^ and *Atg5*^*flox/flox*^ mice were treated with 50 ng/ml PMA, 750 ng/ml ionomycin (both from Sigma-Aldrich), and GolgiPlug (BD Biosciences) in complete medium at 37 °C for 4-6 h. Cells were fixed and permeabilized with TF Fixation/Permeabilization solution (Invitrogen) before interferon gamma (IFN-γ)-APC (Invitrogen, 17-7311-81, 1:160 dilution) staining, and Foxp3-PE-Cy7 (Invitrogen, 25-5773-82, 1:80 dilution) staining, respectively. LIVE/DEAD™ Fixable Aqua Dead Cell Stain Kit (Invitrogen, L34957) was used to distinguish live and dead cells. Dead cells and doublets were excluded from all analysis. Multiparameter analysis was performed on Attune NxT Acoustic flow cytometer (Invitrogen) and analyzed with the FCS express 7 software (De Novo Software, Glendale, CA).

### CD4^+^ T cell isolation and *in vitro* iTreg differentiation

Naïve mouse CD4^+^ T cells were isolated from spleens of *Atg5*^*+/+*^ and *Atg5*^*flox/flox*^ mice using Naïve CD4^+^ T Cell Isolation Kit (Miltenyi Biotec, 130-104-453). Cells (5 × 10^5^ cells in one milliliter per well) were seeded in 24-well plate pre-coated with 0.05 mg/ml goat anti-hamster antibody (MP Biomedicals) at 4°C overnight. iTreg induction medium is glutamine-free RPMI 1640 supplemented with 10% fetal bovine serum (FBS, Gibco), 2-mercaptoethanol (50 µM), penicillin (100 U/ml), streptomycin (100 µg/ml, Corning), hamster anti-CD3 (0.25 µg/ml, eBioscience, 145-2C11), hamster anti-CD28 (1 µg/ml, eBioscience, 37.51), TGF-β (3 ng/ml, Peprotech), and mIL-2 (100 U/ml, Biolegend). Fresh media with escalating concentrations of glutamine were replenished every day for 3 days. Cells were then stained with viability dye, CD4, CD25, Foxp3, and followed by flow cytometric analysis.

### Tumor tissue and plasma glutamine quantification

Glutamine extraction and quantification from tumor tissue was modified from method by Pan et al. [45]. Approximately 50 mg of fresh tumor tissues were homogenized in ice cold 70% ethanol using TissueLyser II (Qiagen). After spinning down, the pellet was collected and dried using SpeedVac vacuum concentrator. The dried pellet was then resuspended in phosphate buffer (20 mM, pH 7.2, 500 µl) and centrifuged to remove debris. The supernatant (400 µl) was transferred and dried with speed vacuum. Subsequently, the dried pellet was dissolved in 550 µl D_2_O containing 0.01 mg/ml Sodium 2,2-Dimethyl-2-Silapentane-5-Sulfonate (DSS; Cambridge Isotope). The samples were then vortexed and centrifuged at 16000 rpm for 12 min. 500 µl solution was transferred to NMR tube. The NMR experiments were carried out at 25°C on a Bruker 700 MHz Avance spectrometer equipped with cryoprobe. To suppress residual macromolecule signals, a Carr-Purcell-Meiboom-Gill (CPMG) sequence with Periodic Refocusing Of J Evolution by Coherence Transfer (PROJECT) method [46] and pre-saturation was used to acquire the 1D ^1^H data. The spectrum width is 13.4 ppm, the recycle delay, acquisition time are 1.5 and 3.5 seconds, respectively. The CPMG duration is 250 ms with 52 echoes and 1.2 ms delay between pulses in the CPMG echo. A control sample with known concentration of glutamine and glutamate and internal reference DSS was prepared to determine the CPMG effect on the peak intensity of glutamine and glutamate. The glutamine and glutamate concentration from tissue extractions was first determined using Chenomx software, and then adjusted by taking account of CPMG effect. Glutamine concentration of the plasma samples was measured by EnzyChrom Glutamine Assay Kit (BioAssay Systems, EGLN-100) following the protocol of the manufacturer. Mouse plasma was collected and diluted two-fold in PBS. To 30 µl of diluted plasma, 15 µl of inactivation solution (0.6 N HCl) was added, mixed, and incubated for 5-10 min at room temperature. Then 15 µl of Tris solution (600 mM, pH 8.5) was added and proceeded with the EnzyChrom Assay.

### αKG extraction and measurement

Alpha-ketoglutarate (αKG) extraction from tumor tissue was as described above. α-ketoglutarate was quantified using αKG Assay Kit (Abcam, ab83431) following the manufacture’s protocol.

## Results

### Attenuated autophagy is sufficient for the suppression of MSTs

To investigate the role of autophagy in regulating MST progression at various stages, we developed an inducible *KRAS*^*G12V*^*;Atg5*^*flox/flox*^ mouse model with an ability for conditional activation of oncogenic KRAS^G12V^ and disruption of the essential autophagy protein ATG5 in submandibular glands (SMGs) [40-42]. We showed that ATG5-knockout tumors grow more slowly during late tumorigenesis, despite a faster onset [40]. MST cells were isolated from both *KRAS*^*G12V*^*;Atg5*^*+/+*^ and *KRAS*^*G12V*^*;Atg5*^*Δ/Δ*^ tumors, respectively for biochemical analyses (**Fig. S1A**). In *KRAS*^*G12V*^*;Atg5*^*Δ/Δ*^ tumor cells, the *Atg5* expression was ablated and the conversion of microtubule-associated protein 1A/1B-light chain 3 (LC3)-I to the lipidated form of LC3B-II was lower than *KRAS*^*G12V*^*;Atg5*^*+/+*^ MSC cells (**Fig. S1B**), supporting the deletion of ATG5. Next, to investigate the effect of host *Atg5* genotype on autophagy competency, we compared autophagy parameters between SMGs and spleens from naïve *Atg5*^*+/+*^ and *Atg5*^*flox/flox*^ mice. As shown in **Fig. 1A**, a reduction, but not depletion, of ATG5-ATG12 and an accumulation of LC3-I confirmed the attenuated autophagy in SMGs and spleens from naïve *Atg5*^*flox/flox*^ mice. Autophagy plays a crucial role in modulating immune system homeostasis [21-25]. Several key autophagy components participate in the immune and inflammatory processes; more specifically, the ATG5-ATG12 conjugate is associated with innate antiviral immune responses [47, 48]. Consistent with this, the basal expression of proinflammatory cytokine genes, *Il-6, Il-1α, Il-1β, Tnfα, Ifn*-γ and *p21*, was significantly higher in SMGs of naïve *Atg5*^*flox/flox*^ mice when compared to SMGs of naïve *Atg5*^*+/+*^ mice (**Fig. 1B**).

**Fig. 1.**
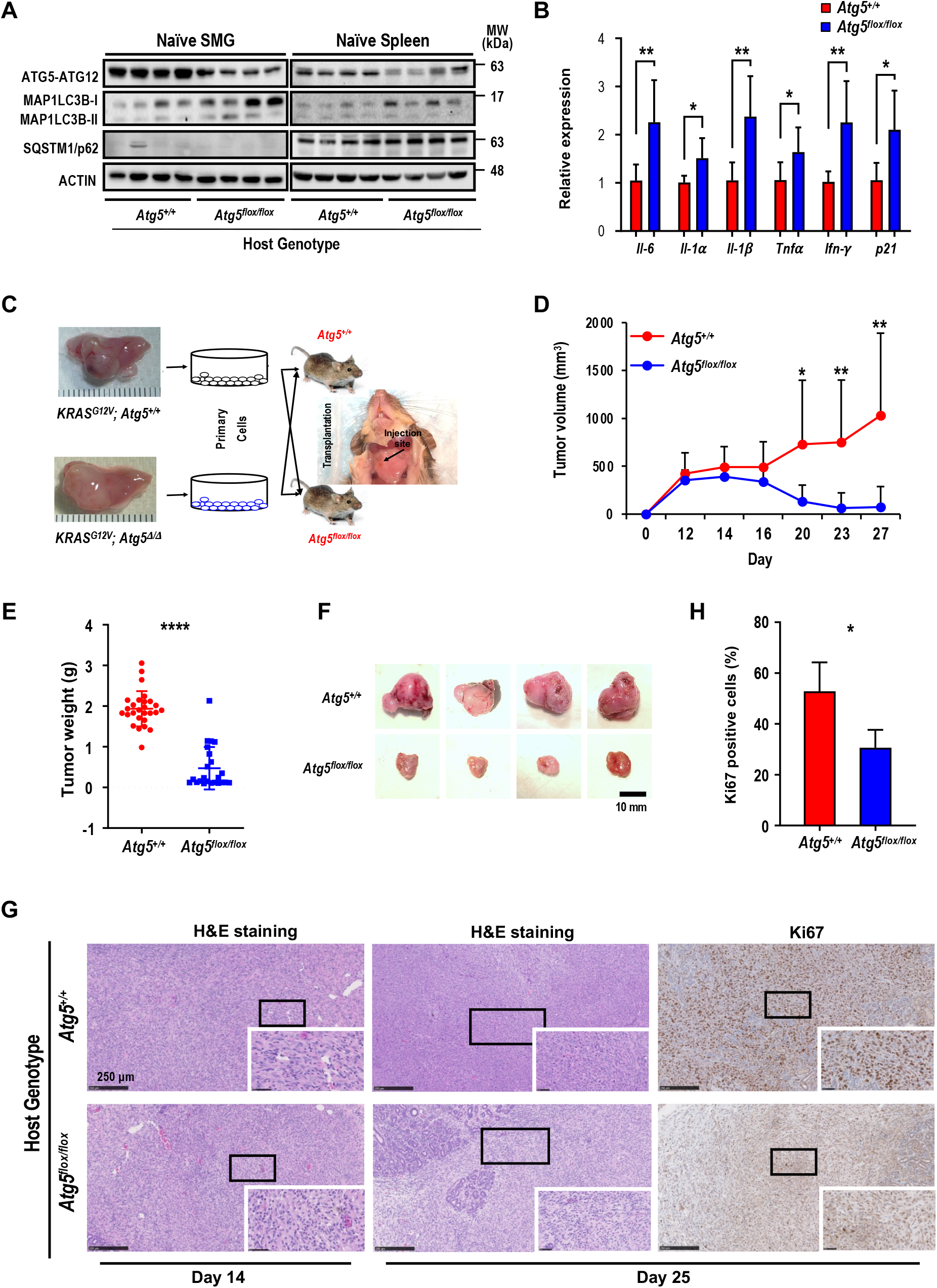
Attenuated autophagy is essential for the suppression of malignant salivary tumors. (**A**) Autophagy activity was verified in SMGs and spleens from *Atg5*^*flox/flox*^ mice by determining the expression of ATG5 and decreased ratio of LC3-II/I. A representative Western blot analysis of ATG5 and basal LC3 in SMGs and spleens from *Atg5*^*+/+*^ and *Atg5*^*flox/flox*^ mice following salivary tumor cell inoculation. (**B**) Quantitative RT-PCR analyses show basal expression of selected proinflammatory cytokine genes in SMGs from naïve *Atg5*^*+/+*^ (n = 3) and *Atg5*^*flox/flox*^ (n = 5) mice. (**C**) Schematic diagram of orthotopic allograft of salivary tumor cells in right SMGs. Host genotypes are designated as *Atg5*^*+/+*^ and *Atg5*^*flox/flox*^, while the injected tumor cell genotypes are designated as *KRAS*^*G12V*^*;Atg5*^*+/+*^ and *KRAS*^*G12V*^*;Atg5*^*Δ/Δ*^. (**D, E, F**) Compromised host autophagy reduces orthotopically implanted salivary tumor expansion. (**D**) Tumor volumes were recorded at 2-day intervals. A representative tumor growth curve is shown. The limitation in tumor growth in host recipients with ATG5 deficiency was observed starting from Day 16 following salivary tumor cell inoculation. (**E, F**) Tumor-bearing SMG weights were measured, and images were taken at Day 25 post-tumor cell inoculation or at humane endpoints. *Atg5*^*+/+*^ and *Atg5*^*flox/flox*^ mice were injected with 2 × 10^5^ primary tumor cells (*KRAS*^*G12V*^*;Atg5*^*+/+*^ and *KRAS*^*G12V*^*;Atg5*^*Δ/Δ*^) in right submandibular glands of *Atg5*^*+/+*^ (n = 18) and *Atg5*^*flox/flox*^ (n = 15) mice (**E**). Representative images of salivary tumors harvested from of *Atg5*^*+/+*^ and *Atg5*^*flox/flox*^ mice (**F**). (**G**) Hematoxylin and eosin (H&E) staining and Ki-67 immunohistochemical staining of SMG tumors. At Day 14 and Day 25 post-implantation, SMGs tissue samples from *Atg5*^*+/+*^ and *Atg5*^*flox/flox*^ mice were collected, processed, and stained with H&E and an anti-Ki-67 antibody for IHC. (**H**) Quantification of Ki-67^+^ cells is as shown (*Atg5*^*+/+*^: n = 6; *Atg5*^*flox/flox*^: n = 4). Five random low-power fields were quantified from each mouse. Scale bar, 250 µm and 50 µm (enlarged view); respectively. Data are shown as mean ± SD; *: *p* < 0.05; **: *p* < 0.01; ****: *p* < 0.0001; Student’s *t*-test, 2-tailed, unpaired.

Next, we hypothesized that an attenuated host autophagy is a barrier for tumor progression in SMGs. To test this possibility, we investigated host tumor microenvironment following the inoculation of *KRAS*^*G12V*^*;Atg5*^*+/+*^ and *KRAS*^*G12V*^*;Atg5*^*Δ/Δ*^ tumor cells, respectively, into different genotypic recipient mice (*Atg5*^*+/+*^ and *Atg5*^*flox/flox*^; **Fig. 1C**). There was no noticeable difference in tumor expansion between *KRAS*^*G12V*^*;Atg5*^*+/+*^ and *KRAS*^*G12V*^*;Atg5*^*Δ/Δ*^ tumor cells in host with the same genotype (lane 1 vs lane 2, lane 3 vs lane 4; **Fig. S1C**). We therefore chose to use *KRAS*^*G12V*^*;Atg5*^*Δ/Δ*^ tumor cells in the subsequent studies for consistency. Tumors derived from the inoculated MST cells exhibited similar histopathological features as the endogenous tumors (**Fig. S1D**). In contrast, a significant reduction in tumor growth, starting from Day 16 following tumor cell inoculation was observed in *Atg5*^*flox/flox*^ recipient mice, compared to *Atg5*^*+/+*^ recipient mice (**Fig. 1D**). Accordingly, SMG tumor weights from *Atg5*^*flox/flox*^ recipient mice were significantly lower than those from *Atg5*^*+/+*^ recipient mice at the later time-point when tumors were harvested (**Fig. 1E, F, SIC**).

H&E staining shows that tumors from *Atg5*^*flox/flox*^ recipient mice often exhibited reduced progression as normal salivary tissues were abundantly detected within SMGs, whereas the SMGs from *Atg5*^*+/+*^ recipient mice had fewer regions displaying normal salivary tissues at Day 14 and Day 25 post-MST cell inoculation (**Fig. 1G**). Consistently, a decrease in the proliferation marker, Ki-67, was observed in SMGs from tumor-bearing *Atg5*^*flox/flox*^ mice compared to those SMG from tumor-bearing *Atg5*^*+/+*^ mice (**Fig. 1G, H**). Of note, tumor growth prior to Day 16 was indistinguishable between two host genotypes (**Fig. 1D**). However, tumor volume between recipient hosts consistently diverged after Day 16. At this point, continued growth was noted only in tumor-bearing *Atg5*^*+/+*^ mice. In contrast, tumor regression was seen in tumor-bearing *Atg5*^*flox/flox*^ mice. These findings underscore the importance of antitumor immune response in a host autophagy-specific manner. Subsequent studies were focused on analyzing the TME residents in tumor-bearing *Atg5*^*+/+*^ mice and *Atg5*^*flox/flox*^ mice on Day 14 and on Day 25, respectively.

### Attenuated autophagy suppresses macrophages expansion within TME

Next, we hypothesized that attenuated autophagy promotes antitumor immunity by affecting the infiltrated cell populations in SMGs. To test this hypothesis, we sought to examine whether autophagy regulates TME residents in our orthotopic syngeneic mouse MST model. Based on our observations of a marked infiltration of inflammatory cells, including macrophages and leukocytes in inducible MSTs [41] and elevated proinflammatory cytokines in naïve *Atg5*^*flox/flox*^ mice (**Fig. 1B**), we first analyzed and compared leukocytes between tumor-bearing *Atg5*^*+/+*^ and *Atg5*^*flox/flox*^ mice. Flow cytometry analyses of CD11b^+^CD49b^+^NK1.1^+^ NK cells (**Fig. S2A**) and CD11b^+^F4/80^-^ Ly6B^+^ neutrophils (**Fig. S2B**) showed that the frequencies of NK cells and neutrophils within the harvested tumor-harboring SMGs at Day 14 post-tumor cell implantation were not host genotype-dependent (**Fig. S2C, D**). At Day 25, NK cell frequency remained the same between different hosts (**Fig. S2E**). However, there was a significant decrease in neutrophils in spleens, a major secondary organ in the immune system, but not SMGs of tumor-bearing *Atg5*^*flox/flox*^ mice at Day 25 (**Fig. S2F**).

We next characterized tumor-associated macrophages (TAMs). Circulating monocytes give rise to mature macrophages that are recruited into the TME and differentiate *in situ* into TAMs upon activation [49]. TAMs are further classified into classically activated or pro-inflammatory M1 and alternatively activated or anti-inflammatory M2 macrophages [50]. We evaluated the correlation between the autophagy capacity and the abundance of macrophage F4/80 marker in SMGs of *Atg5*^*+/+*^ and *Atg5*^*flox/flox*^ recipient mice using immunohistochemistry (IHC). As shown in **Fig. 2A**, a decrease in infiltrating F4/80 counts in SMGs of tumor-bearing *Atg5*^*flox/flox*^ mice at both Day 14 and Day 25 was noted. Flow cytometry analyses further revealed that the percentages of CD11b^+^F4/80^+^MHCII^+^CD86^+^ M1 and CD11b^+^F4/80^+^MHCII^-^ CD206^+^ M2 macrophages in SMGs and spleens (controls) at Day 14 were not host-dependent (**Fig. 2B, C**). Given that tumor burden at Day 25 was higher in SMGs of *Atg5*^*+/+*^ mice than *Atg5*^*flox/flox*^ counterparts (**Fig. 1D**), *Atg5*^*+/+*^ SMGs had increased M1 and M2 macrophage infiltrate compared to *Atg5*^*flox/flox*^ SMGs at Day 25 (**Fig. 2D**). Further, M1 and M2 populations were significantly higher in spleens of tumor-bearing *Atg5*^*+/+*^ and *Atg5*^*flox/flox*^ mice at Day 25 (**Fig. 2E**). Spleen-derived macrophages were readily polarized into M1 and M2 states, presumably via the tumor-spleen signaling interaction of IFN-γ or other cytokines [51, 52]. Together, our data suggests that autophagy promotes the expansion of both M1 and M2 TAMs in SMGs of tumor-bearing *Atg5*^*+/+*^ recipient mice at the later stage post-tumor cell inoculation.

**Fig. 2.**
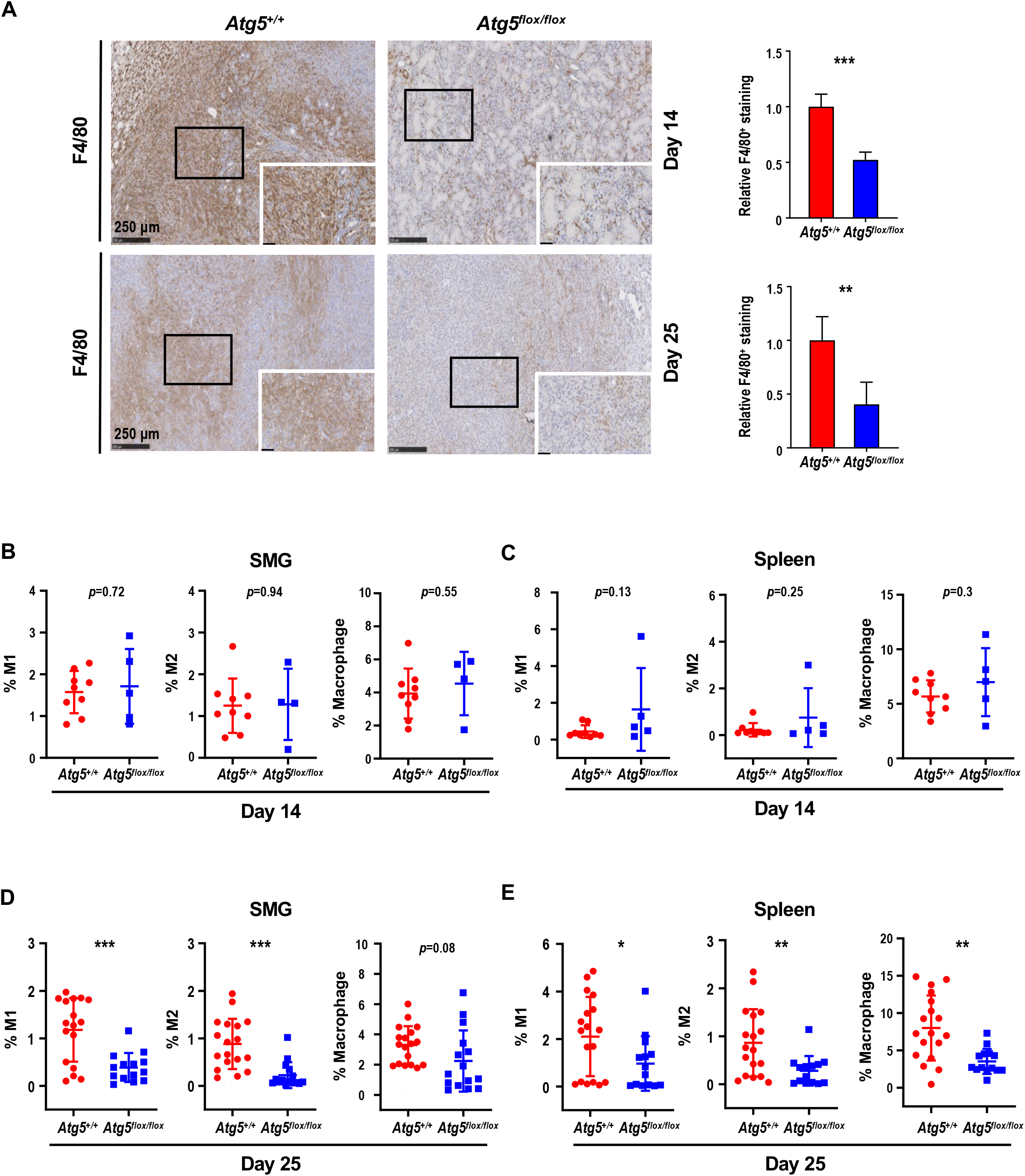
A decrease in M1 and M2 macrophages within TME in tumor-bearing *Atg5*^*flox/flox*^ mice. (**A**) Representative immunohistochemical staining of pan-macrophage marker F4/80 were performed on Day 14 (*upper left two panels*) and Day 25 (*lower left two panels*) in SMG tumors from *Atg5*^*+/+*^ and *Atg5*^*flox/flox*^ mice. Scale bar, 250 µm and 50 µm (enlarged view); respectively. Quantification of F4/80 in Day 14 (*upper right panel*) and Day 25 (*lower right panel*) SMG tumors is as shown. Five random low-power fields were quantified from each mouse. (Day 14, *Atg5*^*+/+*^: n=4 and *Atg5*^*flox/flox*^: n = 4; Day 25, *Atg5*^*+/+*^: n = 6 and *Atg5*^*flox/flox*^: n = 3.) (**B, C**) Flow cytometry analyses show the percentage of M1 macrophages (CD11b^+^F4/80^+^MHCII^+^CD86^+^, *left panel*), M2 macrophages (CD11b^+^F4/80^+^MHCII^-^CD206^+^, *middle panel*) and macrophages (CD11b^+^F4/80^+^, *right panel*) in alive SMG cells (**B**) and splenocytes (**C**) of tumor-bearing mice at Day 14 post-implantation. (**D, E**) Flow cytometry analyses show the percentage of M1 macrophages (CD11b^+^F4/80^+^MHCII^+^CD86^+^, *left panel*), M2 macrophages (CD11b^+^F4/80^+^MHCII^-^CD206^+^, *middle panel*) and macrophages (CD11b^+^F4/80^+^, *right panel*) in alive SMG cells (**D**) and splenocytes (**E**) of tumor-bearing mice at Day 25 post-inoculation. *Atg5*^*+/+*^: n = 18 and *Atg5*^*flox/flox*^: n = 15. Data are shown as mean ± SD; *: *p* < 0.05; **: *p* < 0.01; ***: *p* < 0.001; Student’s *t*-test, 2-tailed, unpaired.

### Attenuated autophagy promotes tumor-infiltrating cytotoxic CD8^+^ T cells and IFN-γ production

Cancer immune evasion is a major stumbling block for antitumor immunity. We further elucidated the role of autophagy in regulating the infiltration of cytotoxic T lymphocytes into tumors. To achieve this goal, we first focused on CD4^+^ helper and CD8^+^ cytotoxic T lymphocytes. At Day 14 post-tumor cell implantation, there was no significant difference in the frequency of infiltrating CD8^+^ T cells between SMGs of tumor-bearing *Atg5*^*+/+*^ and *Atg5*^*flox/flox*^ mice by IHC (**Figs. 3A**). However, a clear increase in the percentage of infiltrating CD4^+^ and CD8^+^ T cells was detected in SMGs of tumor-bearing *Atg5*^*flox/flox*^ mice at Day 25 by IHC (**Fig. 3A**). Consistently, flow cytometry (**Fig. 3B, C)** confirmed a higher percentage of CD8^+^ T cells in SMGs (**Fig. 3D, F**, *left panels*) and spleens (**Fig. 3E, G**, *left panels*), respectively, in tumor-bearing *Atg5*^*flox/flox*^ mice at both Day 14 (**Fig. 3D, E**) and Day 25 (**Fig. 3F, G**). Conversely, the host autophagy capacity did not affect percentages of CD4^+^ T cells in SMGs and spleens from tumor-bearing mice at Day 14 (**Fig. 3D, E**, *right panels*) and Day 25 (**Fig. 3F, G**, *right panels*). We conclude that the increased infiltrating CD8^+^ cytotoxic T cells play a key role in the observed antitumor phenotypes in tumor-bearing *Atg5*^*flox/flox*^ mice.

**Fig. 3.**
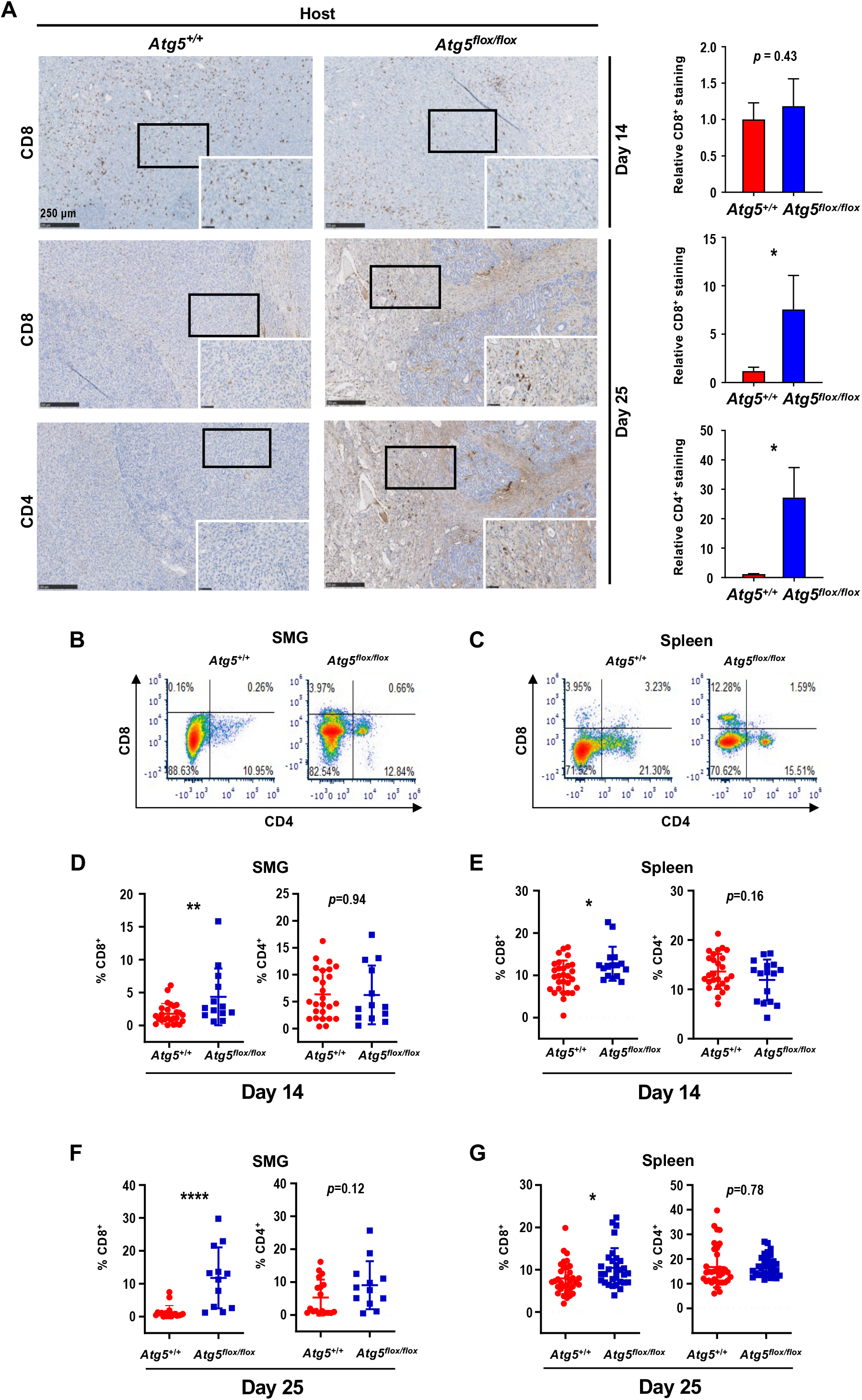
An enhancement in CD8^+^ T cells within TME in tumor-bearing *Atg5*^*flox/flox*^ mice. (**A**) Representative immunohistochemistry analyses of CD8 on Day 14 SMG tumors (*upper left panels*), and CD8 (*middle left panels*) and CD4 (*lower left panels*) on Day 25 SMG tumors from *Atg5*^*+/+*^ and *Atg5*^*flox/flox*^ mice. Scale bar, 250 µm and 50 µm (enlarged view); respectively. Quantification of CD8^+^ signals in Day 14 SMG tumors (*upper right panel*) and quantification of CD8^+^ signals (*middle right panel*) and CD4^+^ signals (*lower right panel*) in SMG tumors from Day 25 *Atg5*^*+/+*^ and *Atg5*^*flox/flox*^ mice. Five random low-power fields were quantified from each mouse. *Atg5*^*+/+*^: n ≥ 4 and *Atg5*^*flox/flox*^: n ≥ 3. (**B, C**) Representative flow cytometry showing CD8^+^ T cells and CD4^+^T cells isolated from alive SMG cells (**B**), and splenocytes (**C**) of tumor-bearing *Atg5*^*+/+*^ and *Atg5*^*flox/flox*^ mice. (**D**-**G**) Flow cytometry analyses showing the percentage of CD8^+^ T cells and CD4^+^ T cells in alive SMG tumor cells, and splenocytes from Day 14 (**D, E**) and Day 25 (**F, G**) tumor-bearing *Atg5*^*+/+*^ (*red*) and *Atg5*^*flox/flox*^ (*blue*) mice (*Atg5*^*+/+*^ and *Atg5*^*flox/flox*^: n ≥ 12). Data are shown as mean ± SD; *: *p* < 0.05; **: *p* < 0.01; ****: *p* < 0.0001; Student’s *t*-test, 2-tailed, unpaired.

IFN-γ is key to cellular immune responses and is secreted predominantly by activated lymphocytes, such as CD4^+^ helper and CD8^+^ cytotoxic T cells [53]. We next assessed expression of proinflammatory cytokine-related genes in SMGs from tumor-bearing recipient mice by qRT-PCR and found increased IFN-γ expression in *Atg5*^*flox/flox*^ recipient mice at both Day 14 and Day 25 (**Figs. 4A, B**). However, the IFN-γ production by splenocytes from the naïve *Atg5*^*+/+*^ and *Atg5*^*flox/flox*^ mice challenged with lipopolysaccharide (LPS), to stimulate the release of proinflammatory cytokines, were comparable (**Fig. 4C**). We conclude that different host autophagy capacity did not alter IFN-γ production from splenocytes upon LPS challenge. Next, we treated isolated SMG resident cells and splenocytes with a leukocyte activation cocktail containing PMA/ionomycin/Golgiplug [54] to promote intracellular cytokines accumulations, and assessed IFN-γ -production by flow cytometry (**Fig. S3A, B**). Notably, there was a significant increase in the frequency of IFN-γ -producing cells in SMGs and spleens of tumor-bearing *Atg5*^*flox/flox*^ mice at Day 25 (**Fig. 4E)**, but not at Day 14 (**Fig. 4D**). Further, no changes in CD8^+^IFN-γ + and CD4^+^IFN-γ + populations in SMGs (**Fig. 4F**), and spleens (**Fig. 4G**) from *Atg5*^*+/+*^ and *Atg5*^*flox/flox*^ recipient mice were observed at Day 14. In contrast, *Atg5*^*flox/flox*^ recipient mice had higher frequencies of CD8^+^IFN-γ + cells in SMGs and spleens, respectively, at Day 25 (**Fig. 4H, I**, *left panels*). CD4^+^IFN-γ ^+^ cells, albeit with reduced frequencies, were also higher in SMGs and spleens of *Atg5*^*flox/flox*^ recipient mice (**Fig. 4H, I**, *right panels*). It is conceivable that the increased IFN-γ production by cytotoxic CD8^+^IFN-γ + cells improved the antitumor responses in SMGs of *Atg5*^*flox/flox*^ recipient mice.

**Fig. 4.**
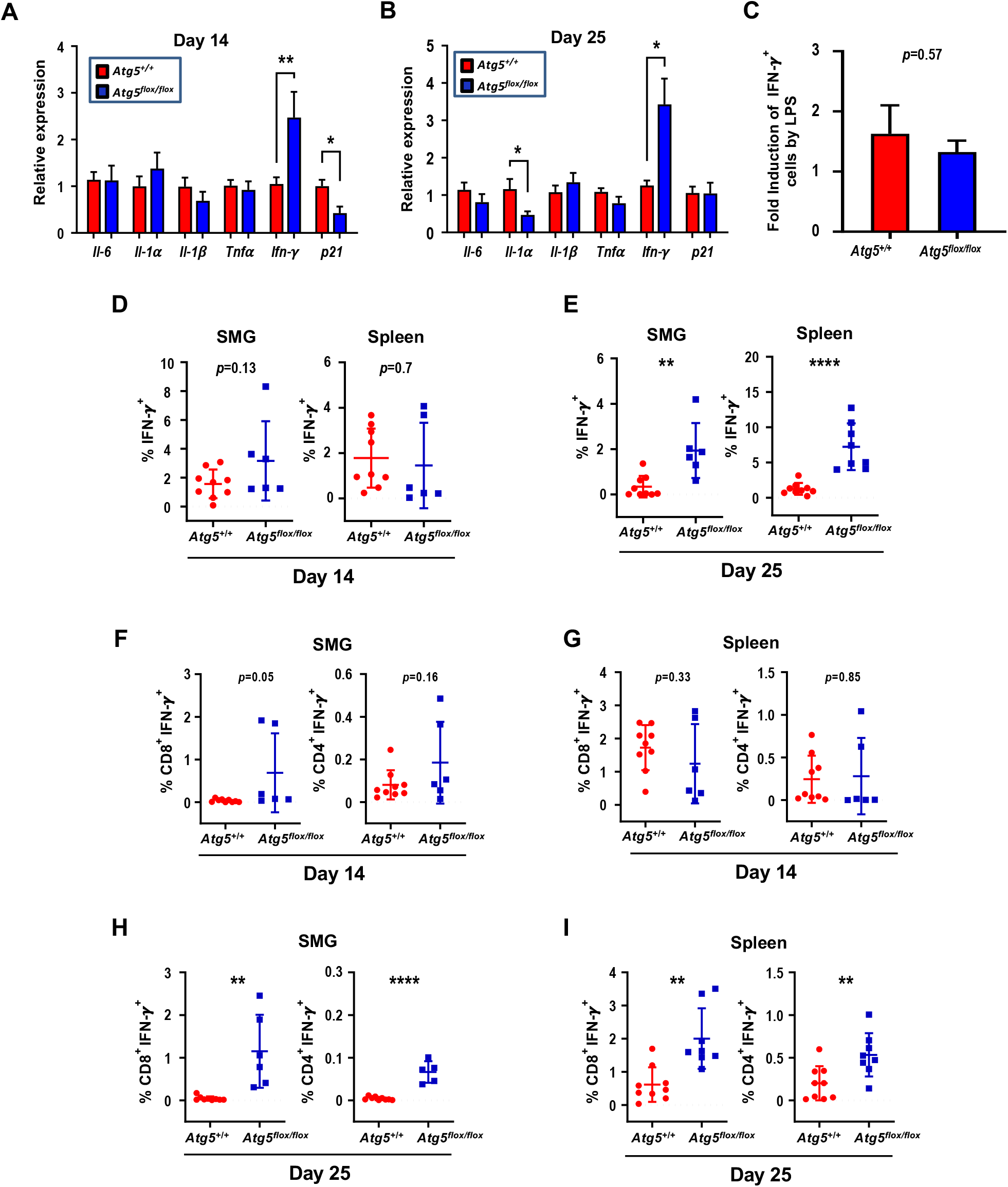
Attenuated autophagy promotes IFN-γ -producing CD8^+^ and CD4^+^ T cells in SMGs and spleens of tumor-bearing mice. (**A**) Relative expression of proinflammatory cytokine-related genes in SMGs from tumor-bearing *Atg5*^*+/+*^ and *Atg5*^*flox/flox*^ mice, Day 14 (**A**) and Day 25 (**B**) post-implantation (n = 3). (**C**) Fold induction of IFN-γ -producing cells in spleens of *Atg5*^*+/+*^ and *Atg5*^*flox/flox*^ mice following LPS stimulation (5 mg/kg) for 6 h compared with that from PBS control mice (n = 3). (**D**) The percentage of IFN-γ -producing cells in single cell suspensions of the alive SMG cells (*left panel*) and splenocytes (*right panel*), from Day 14 tumor-bearing *Atg5*^*+/+*^ (*red*) and *Atg5*^*flox/flox*^ (*blue*) mice, following PMA/ionomycin/Golgiplug stimulation for 4 h. (**E**) The percentage of IFN-γ -producing cells in single cell suspensions of the alive SMG resident cells (*left panel*) and splenocytes (*right panel*), from Day 25 tumor-bearing *Atg5*^*+/+*^ (*red*) and *Atg5*^*flox/flox*^ (*blue*) mice, following PMA/ionomycin/Golgiplug stimulation for 4 h. (**F**-**I**) Flow cytometry analyses showing the percentage of IFN-γ + T cells, CD8^+^IFN-γ + cells and CD4^+^IFN-γ +, in single cell suspensions of the alive SMG cells and splenocytes from Day 14 (**F, G**) and Day 25 (**H, I**) tumor-bearing *Atg5*^*+/+*^ (*red*) and *Atg5*^*flox/flox*^ (*blue*) mice (*Atg5*^*+/+*^ and *Atg5*^*flox/flox*^: n ≥ 5). Data are shown as mean ± SD; *: *p* < 0.05; **: *p* < 0.01; ****: *p* < 0.0001; Student’s *t*-test, 2-tailed, unpaired.

### Glutamine-dependent regulation of Treg cells in SMGs and spleens

Subsets of T cells within TME play distinct roles in mediating antitumor immunity [55]. Upon tumor antigen stimulation, naïve T cells are activated and differentiate into two broad classes of CD4^+^ or CD8^+^ T cells that have distinct effector mechanisms [12]. One CD4^+^ T cell subset, Tregs, dampens the antitumor immune response [18]. We next examined the effect of host autophagy capacity on Treg population within TME of *Atg5*^*+/+*^ and *Atg5*^*flox/flox*^ recipient mice. A lower count of CD4^+^CD25^+^Foxp3^+^ Tregs in SMGs at Day 14 was detected in tumor-bearing *Atg5*^*flox/flox*^ mice (**Fig. 5A**, *left panel*), while the Treg counts in spleens from mice with different host autophagy capacity were indistinguishable (**Fig. 5A**, *right panel*). Notably, **Fig. 5B** showed a significant decrease in Tregs in SMGs and spleens from tumor-bearing *Atg5*^*flox/flox*^ mice at Day 25. During T cell activation, glutamine metabolism increases to meet rapid growth requirement [56]. We have previously shown that autophagy deficiency contributes to reduced intracellular concentration of most amino acids except glutamine, the level of which increased in *KRAS*^*G12V*^*;Atg5*^*Δ/Δ*^ tumor cells [40]. Given that autophagy inhibition promotes glutamine uptake [57], we examined whether glutamine level regulates T cell differentiation into specific subtypes in SMGs of *Atg5*^*+/+*^ and *Atg5*^*flox/flox*^ recipient mice. Notably, both glutamine (**Fig. 5C**, *right panel*) and its metabolite α-ketoglutarate (αKG) (**Fig. 5D**, *right panel*) levels were higher in SMGs from tumor-bearing *Atg5*^*flox/flox*^ mice at Day 14 comparing to tumor-bearing *Atg5*^*+/+*^ mice. In contrast, there was no difference in glutamine and αKG detected in SMGs of naïve *Atg5*^*+/+*^ and *Atg5*^*flox/flox*^ mice (**Fig. 5C, D**, *left panels*). **Fig. 5E** showed that naïve T cells differentiated into Tregs in a reverse glutamine-dependent manner under Treg polarization conditions, suggesting that glutamine shortage would render a higher frequency of CD4^+^CD25^+^Foxp3^+^ Tregs. This finding was consistent with the decreased *Foxp3* expression in Tregs differentiated under increasing glutamine concentrations in polarization medium (**Fig. 5F**). Interestingly, there was an increase in the frequency of IFN-γ secreting CD4^+^ T cells derived from naïve T cells of *Atg5*^*flox/flox*^ mice at 24 h and 48 h in glutamine-replenished Treg polarization medium (**Fig. S4**). Conceivably, glutamine concentration in SMGs not only regulated Tregs population but also IFN-γ secreting CD4^+^ T cells. Together, **Table 1** summarizes the comparison between tumor growth and TME residents from *Atg5*^+/+^ and *Atg5*^*flox/flox*^ recipient mice at Day 14 and Day 25. Tumors continued to grow in the *Atg5*^+/+^ recipient mice while tumors regressed in *Atg5*^*flox/flox*^ recipient mice after 14 days following cell implantation. A statistically significant decrease in CD4^+^ subpopulation, Tregs, and an increase in CD8^+^ T cells were noted (**Table 1**), supporting the role of attenuated host autophagy in promoting antitumor immune responses in MST-bearing *Atg5*^*flox/flox*^ mice.

**Fig. 5.**
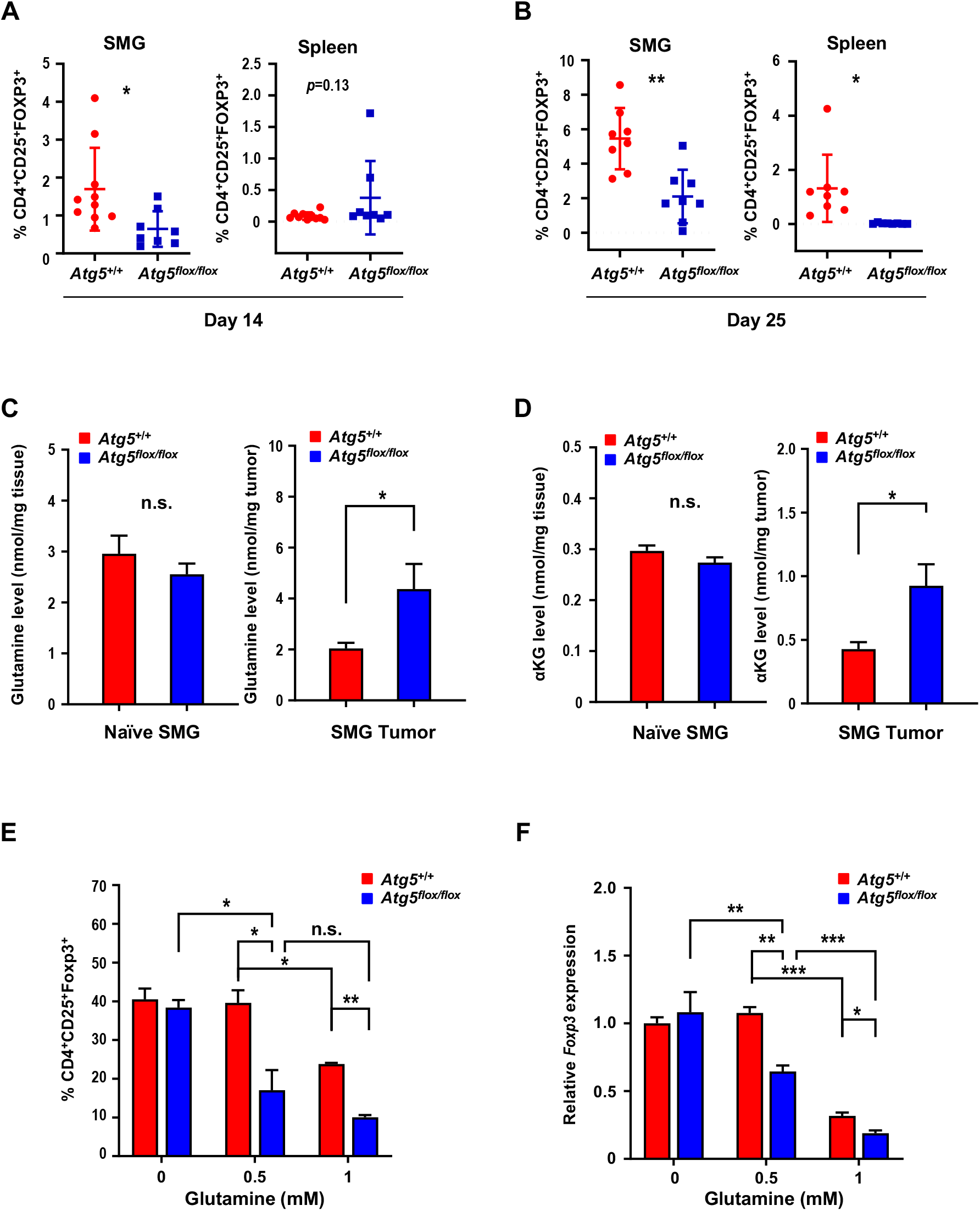
Attenuation of Treg population in SMGs and spleens of *Atg5*^*flox/flox*^ mice is associated with the glutamine concentration in SMG tumor microenvironment. (**A, B**) Flow cytometry analyses showing CD4^+^CD25^+^Foxp3^+^ Tregs in single cell suspensions of the SMG cells (*left panels*) and splenocytes (*right panels*) of tumor-bearing mice, Day 14 (**A**) and Day 25 (**B**) after tumor implantation (n ≥ 8). (**C, D**) Levels of glutamine (**C**) and α-ketoglutarate (αKG; **D**) in naïve SMGs (*left panels*) and Day 14 SMG tumors (*right panels*). Glutamine and αKG concentrations in SMG tumors and SMGs from naïve mice were respectively determined (n ≥ 5). (**E**) The percentage of CD4^+^CD25^+^Foxp3^+^ Tregs, a subset of CD4^+^ T cells, is negatively correlated with glutamine concentration in Treg polarization medium. Naïve mouse CD4^+^ T cells were isolated from mouse spleen and cultured for 3 days in Treg polarization medium with indicated glutamine concentrations. The percentages of Foxp3^+^ cells of total CD4^+^CD25^+^ cells are indicated in the bar graph (n = 3). (**F**) Levels of *Foxp3* mRNA expression in the induced Tregs after cultured in Treg polarization medium with indicated glutamine concentrations for the indicated genotypes. Naïve mouse CD4^+^ T cells were isolated from spleens of *Atg5*^*+/+*^ (*red*) and *Atg5*^*flox/flox*^ (*blue*) mice and cultured 3 days in Treg polarization medium. Expression of Foxp3 mRNA isolated from the differentiated cells was analyzed by qRT-PCR (n = 3). Data are presented in bar graph shown as Mean ± SEM. *p* value was calculated by *t* test (unpaired, two tailed). *: *p* < 0.05; **: *p* < 0.01; ***: *p* < 0.001; *n*.*s*., not significant.

**Table 1:** Immune cell profile in autophagy-deficient versus autophagy-sufficient tumor microenvironments at two selected endpoints, Day 14 and Day 25, post-tumor implantation. Data summarized is based on flow cytometry analyses from **Figs. 2-5** and **S2**.

| Host genotypes | *Atg5<sup>flox/flox</sup>* versus *Atg5<sup>+/+</sup>* |  |
| --- | --- | --- |
| Time points | Day 14 | Day 25 |
| NK cells | – | – |
| Neutrophils | – | – |
| CD4<sup>+</sup> T cells | – | – |
| CD8<sup>+</sup> T cells | ↑ | ↑ |
| CD4<sup>+</sup>IFN-γ<sup>+</sup> | – | ↑ |
| CD8<sup>+</sup>IFN-γ<sup>+</sup> | – | ↑ |
| TAMs (M1 and M2) | – | ↓ |
| Tregs | ↓ | ↓ |

### Dietary glutamine supplementation is sufficient to suppress MST

Next, we evaluated the effect of dietary glutamine supplementation on MST tumor growth. We used isocaloric diet with 20% additional glutamine (high glutamine diet) compared to control diet as reported by Ishak Gabra [43]. Supplementation of glutamine in the diet significantly increases the plasma concentration of glutamine in naïve *Atg5*^+/+^ mice (**Fig. 6A**). Furthermore, comparing to the *Atg5*^+/+^ mice fed with control diet, tumor glutamine level was elevated in the *Atg5*^+/+^ mice fed with high glutamine diet (**Fig. 6B**). These immunocompetent *Atg5*^+/+^ mice fed with high glutamine diet developed significantly smaller tumors after orthotopic tumor implantation (**Fig. 6C**). H&E staining of the excised tumors from mice fed with high glutamine diet showed areas of residual normal glandular parenchyma (**Fig. S5)**. IHC staining revealed that infiltrating cytotoxic CD8^+^ T cells were more notably abundant with overall less Foxp3^+^ Tregs detected in tumor sections from high glutamine-fed mice (**Fig. 6D**, *upper 4 panels*). Notably, tumor PD-LI signals were moderately strong without clear spatial distribution or affected by high glutamine diet (**Fig. 6D**, *lower 4 panels*). Presumably, the increase in tumor-infiltrating CD8^+^ T cells is caused by the reduction of Tregs. Altogether, dietary intake of glutamine may effectively increase the concentration of glutamine in TME to suppress Treg differentiation, mimicking an autophagy-compromised TME (**Table 1**).

**Fig. 6.**
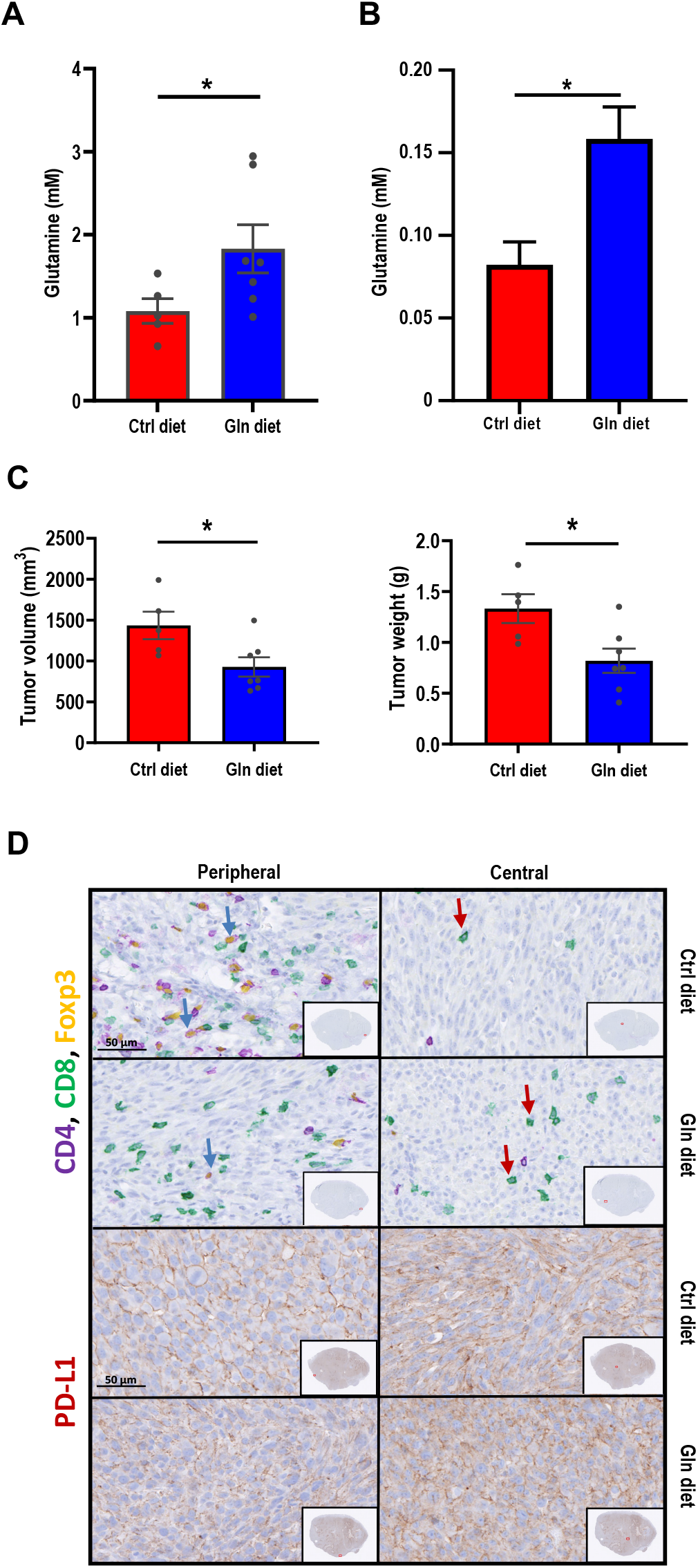
Dietary glutamine supplementation reduces MST tumor burden with increased CD8^+^ cell infiltration. (**A, B**) 28 days of dietary glutamine supplementation increases glutamine concentration in plasma of naïve mice (**A**) and in SMG of tumor-bearing mice (**B**) (n ≥ 5 in each cohort). (**C, D**) Mice with orthotopic MST implantation were fed with control (Ctrl diet, n = 5) and glutamine-supplemented diet (Gln diet, n = 7), respectively, for 21 days prior to tumor implantation and for another 21 days after implantation, prior to euthanasia. Tumor volume (*left*) and wet SMG weight (*right*) at Day 21 post-tumor implantation are shown (**C**), and representative micrographs of indicated IHC stains of the orthotopic MST tumors are shown (**D**). (*Upper 4 panels*) IHC staining for CD4, CD8 and Foxp3 cells in FFPE tumor sections. Green arrows indicate Foxp3-positive (yellow nuclei staining) cells. Red arrows indicate CD8^+^-positive (green membrane staining) cells. Inlets are a low-power overview (red boxes indicate relative spatial location of enlarged views (scale bar: 50 μm). Peripheral (*left*) is considered < 500 μm from the edge, central/core (non-necrotic region) > 500 μm from the edge. (*Lower 4 panels*) IHC staining of tumor sections using PD-L1 antibody (red membrane staining). Images are representatives of ≥ 5 biological replicates. Nuclei were stained with hematoxylin (blue). Data are shown as mean ± SD; *: *p* < 0.05 by Welch *t* test.

## Discussion

Immune suppression and escape are increasingly recognized as critical traits of malignancy [58]. During cancer progression, autophagy may represent an important pathway for immune escape, while also promoting the malignant phenotype of cancer cells [59, 60]. The increasing interests in the role of compromised autophagy, in addition to undermining tumorigenesis, in controlling immune tolerance and bolstering tumor rejection [61], prompted us to develop a syngeneic orthotopic mouse tumor model for evaluating host autophagy capacity on MST progression. This novel syngeneic tumor model enables us to show for the first time that host ATG5-dependent autophagy promotes tumor progression by suppressing the antitumor immune response, independently of the autophagy genotypes of donor tumor cells. In other words, the attenuated host autophagy capacity ultimately results in spontaneous tumor regression and improved survival of tumor-bearing mice through an “antitumor” TME. Autophagy plays a key role in the function and development of neutrophils, macrophages, NK cells, T cells and B cells and dendritic cells [62], key components of TME. In general, the relationship between autophagy and immune system is complex, and there is no consensus on the role autophagy plays in antitumor immunity. Our data suggest at least two of these populations, T lymphocytes and macrophages, are affected by attenuated host autophagy within TME. The improved antitumor TME in *Atg5*^*flox/flox*^ mice is consistent with reports from recent studies that implicate autophagy in immune evasion which may restrain antitumor immunity [63, 64]. For example, Cunha et al. reported that the growth of subcutaneously engrafted murine melanoma is suppressed in ATG5-compromised mice by M1-polarized TAMs and increased type I IFN production [65]. Further, autophagy promotes tumor immune tolerance by enabling Treg function and limiting expression of IFN and CD8^+^ T cell response which in turn enables tumor growth [66]. Likewise, blocking hypoxia-induced autophagy in tumors restores cytotoxic T cell activity and promotes regression in lung cancer [36]. Additionally, loss of host autophagy increases the level of circulating pro-inflammatory cytokines and promotes T cell infiltration in tumors with high tumor mutational burden [66]. Consistent with these reports, we showed that autophagy is a critical immune-suppressing factor that regulates the infiltration and activity of cytotoxic CD8^+^ T cells. We found that attenuation, even not complete depletion, of ATG5 abundance alone is sufficient to increase IFN-γ expression by IFN-γ producing cells at both early and later tumor stages in *Atg5*^*flox/flox*^ mice. Tumor-infiltrating CD8^+^ T cells is a useful prognostic parameter in various cancers [67]. Indeed, CD8^+^ T cells are restrained due to long-lasting interactions with TAMs, whereas depletion of TAMs restores T cell migration and infiltration into tumor islets [68]. Consistently, we found that there are more macrophages infiltrating the tumors in *Atg5*^+/+^ mice, which may impede migration of CD8^+^ T cells into the TME.

Herein, we also report glutamine to be an immunometabolic regulator in SMGs that links compromised autophagy to immunosuppressive Foxp3^+^ Tregs. Tregs play a crucial role in the prevention of antitumor immunity by suppressing the activation and differentiation of CD4^+^ helper T cells and CD8^+^ cytotoxic T lymphocytes [69]. In this study, we found that increased Tregs infiltrate was accompanied with low CD8^+^IFN-γ + infiltrate in SMGs and spleens in tumor-bearing *Atg5*^+/+^ mice at Day 25. Higher Treg infiltrate within SMG TME of tumor-bearing *Atg5*^+/+^ mice would inhibit CD8^+^ cytotoxic T cells, leading to a tumor progression phenotype. Moreover, we found that glutamine supplementation inhibited the skewing of naïve T cells isolated from the spleen into Tregs. In concordance with our previous studies [40], the difference in Intratumoral glutamine level was prominent between SMGs of *Atg5*^*+/+*^ and *Atg5*^*flox/flox*^ recipient mice (**Fig. 5C**). Glutamine is a non-essential, but the most abundant amino acid in the body [70]. It participates in central metabolic processes by acting as an energy substrate for the tricarboxylic acid cycle and a nitrogen donor in several pathways including purine/pyrimidine synthesis, nicotinamide adenine dinucleotide metabolism, and the urea cycle [71, 72]. Several mechanisms have been suggested to link glutamine and antitumor immunity. In macrophages and T cells, it has been reported to be mediated via shifts in energy utilization (*i*.*e*., the balance between glycolysis and glutaminolysis), which alters the levels of intermediary metabolites such as αKG [73]. A report by Tran et al. showed that glutamine-αKG axis suppresses Wnt signaling and promotes cellular differentiation, thereby restricting tumor growth in colorectal cancer [74]. Furthermore, αKG-dependent demethylation is a critical regulatory step in T cell activation and differentiation and macrophage polarization [75-77]. Herein, we demonstrate that high glutamine levels reduced CD4^+^CD25^+^Foxp3^+^ Treg cell population (**Fig. 5E, F**). Here, we established a causal link between dietary glutamine supplementation and antitumor immunity in mouse MST was established.

Nutritional stress is used by cancer cells to generate an immunosuppressive microenvironment to impact the function of tumor-infiltrating lymphocytes [13, 78-80]. Within tumors, intratumor nutrient level is determined by the net balance of host blood supply, autophagy, and the net competition between tumor cells and other TME residents [78]. In addition, the increased metabolic demands of tumor cells and activated T lymphocytes may introduce competition for glutamine within the TME [81], creating a scenario in which tumor cells out-compete T cells for local glutamine and thereby alter the characteristics of the tumor-infiltrating lymphocytes. Thus, in this scenario the glutamine consumption would both promote proliferation and survival of tumor cells and limit the capacity for T cell-mediated antitumor immunity simultaneously, similar to observations with arginine [78]. Accordingly, we postulated that glutamine-consuming tumors, such as MSTs, might benefit therapeutically from dietary glutamine supplementation, improving antitumor T cell responses by reversing a tumor “glutamine grab” phenomenon. However, other immune cells may also be affected by dietary glutamine supplementation. For example, the production of αKG via glutaminolysis is important for activation of M2-like macrophages [77]. Thus, although our findings of improved intratumoral T cell effector functions likely result from the increased glutamine availability to suppress Tregs, the potential remains for additional factors that can impact the immune system such as inhibition of suppressive microenvironments by M2-like macrophages. It is also possible that different residents within TME use distinct nutrients according to their own unique metabolic programs. The dietary glutamine supplementation enables a metabolic signaling pathway that suppress the function of some immune system T cells to promote others. The concept of preventing the “glutamine steal” by tumor cells as a treatment strategy may be applicable beyond MSTs as multiple types of tumors are also considered to be glutamine-addicted. It is possible that this phenomenon is playing out in other cancers as well.

## Conclusions

In summary, we found an elevated expression of basal proinflammatory cytokines in the SMGs of naïve *Atg5*^*flox/flox*^ mice with attenuated autophagy. Subsequently, we developed a syngeneic orthotopic MST tumor model in *Atg5*^*+/+*^ and *Atg5*^*flox/flox*^ mice and revealed that *Atg5*^*flox/flox*^ mice suppressed orthotopically allografted MST cells. Together with reduced growth of tumors, there was an enhanced antitumor immune response demonstrated by reduction of both M1 and M2 macrophages, increased infiltration of CD8^+^ T cells, elevated IFN-γ production, as well as decreased inhibitory Tregs within TME and spleens of recipient mice. Mechanistically, attenuated autophagy led to increased levels of glutamine within SMGs which in turn would promote the inflammatory T cells while inhibiting the generation of Tregs in tumor-bearing *Atg5*^*flox/flox*^ mice. In addition, dietary glutamine supplementation, mimicking attenuated autophagy, retarded tumor expansion in *Atg5*^*+/+*^ mice.

## Abbreviations

αKG: α-ketoglutarate
ATG3: autophagy-related 3
ATG5: autophagy-related 5
ATG7: autophagy-related 7
ATG12: autophagy-related 12
CD25: cluster of differentiation 25
CD28: cluster of differentiation 28
CD3: cluster of differentiation 3
CD4: cluster of differentiation 4
CD8: cluster of differentiation 8
FOXP3: forkhead box P3
IFN-γ: interferon gamma
IL1: interleukin-1
IL2: interleukin-2
IL6: interleukin-6
Treg: regulatory T cell
KRAS: kirsten rat sarcoma viral oncogene homolog
LPS: lipopolysaccharide
MST: malignant salivary gland tumor
NK: natural killer cell
P21: cyclin dependent kinase inhibitor 1A
SDC: salivary duct carcinoma
SMG: submandibular gland
TAM: tumor-associated macrophage
TGFB: transforming growth factor beta
Th1: type 1 T helper cell
Th2: type 2 T helper cell
TME: tumor microenvironment
TNFA: tumor necrosis factor alpha
Treg: regulatory T cell.

## Acknowledgements

We thank members of Dr. Ann’s laboratory for helpful discussions on the manuscript. This work was supported in part by funds from the National Institutes of Health R01DE10742, R21DE023298 and R01DE026304 (to D.K.A.), and P30CA033572 (supporting research work carried out in Core Facilities).

## Author Contributions

S.C., Y.C., Y.-C.W., Y.-W.H., Y.Q., X.Z., H.H.L., and D.K.A. designed the experiments and analyzed the data. S.C., Y.C., Y.-C.W., Y.-W.H., Y.Q., W.H., and A.C. executed experiments. C.O. analyzed public datasets. S.C., H.H.L., Z.S., E.M. and D.K.A. prepared the manuscript. All authors have commented on the manuscript.

