## Supplementary Figures and Table for "Glutamine is essential for overcoming the immunosuppressive microenvironment in malignant salivary gland tumors"

**Shuting Cao[a], Yu-Wen Hung[a, #], Yi-Chang Wang[a, #], Yiyin Chung[a, #], Yue Qi [a, #], Ching Ouyang[b], Xiancai Zhong[c], Weidong Hu[c], Alaysia Coblentz[a], Ellie Maghami[d], Zuoming Sun[c, e], H. Helen Lin[a], and David K. Ann[a, e, *]**

[a]Department of Diabetes Complications and Metabolism, Arthur Riggs Diabetes and Metabolism Research Institute, Beckman Research Institute, City of Hope, Duarte, CA, 91010, USA

[b]Department of Computational and Quantitative Medicine, Beckman Research Institute, City of Hope Comprehensive Cancer Center, Duarte, CA 91010, USA

[c]Department of Immunology and Theranostics, Beckman Research Institute, City of Hope Comprehensive Cancer Center, Duarte, CA 91010, USA

[d]Division of Head and Neck Surgery, City of Hope National Medical Center, Duarte, CA 91010, USA

[e]Irell & Manella Graduate School of Biological Sciences, Beckman Research Institute, City of Hope Comprehensive Cancer Center, Duarte, CA 91010, USA

**[#]Contribute equally**
***To whom all correspondence should be addressed:**
David K. Ann, Ph.D.
Beckman Research Institute
City of Hope Comprehensive Cancer Center
Duarte, CA 91010-3000

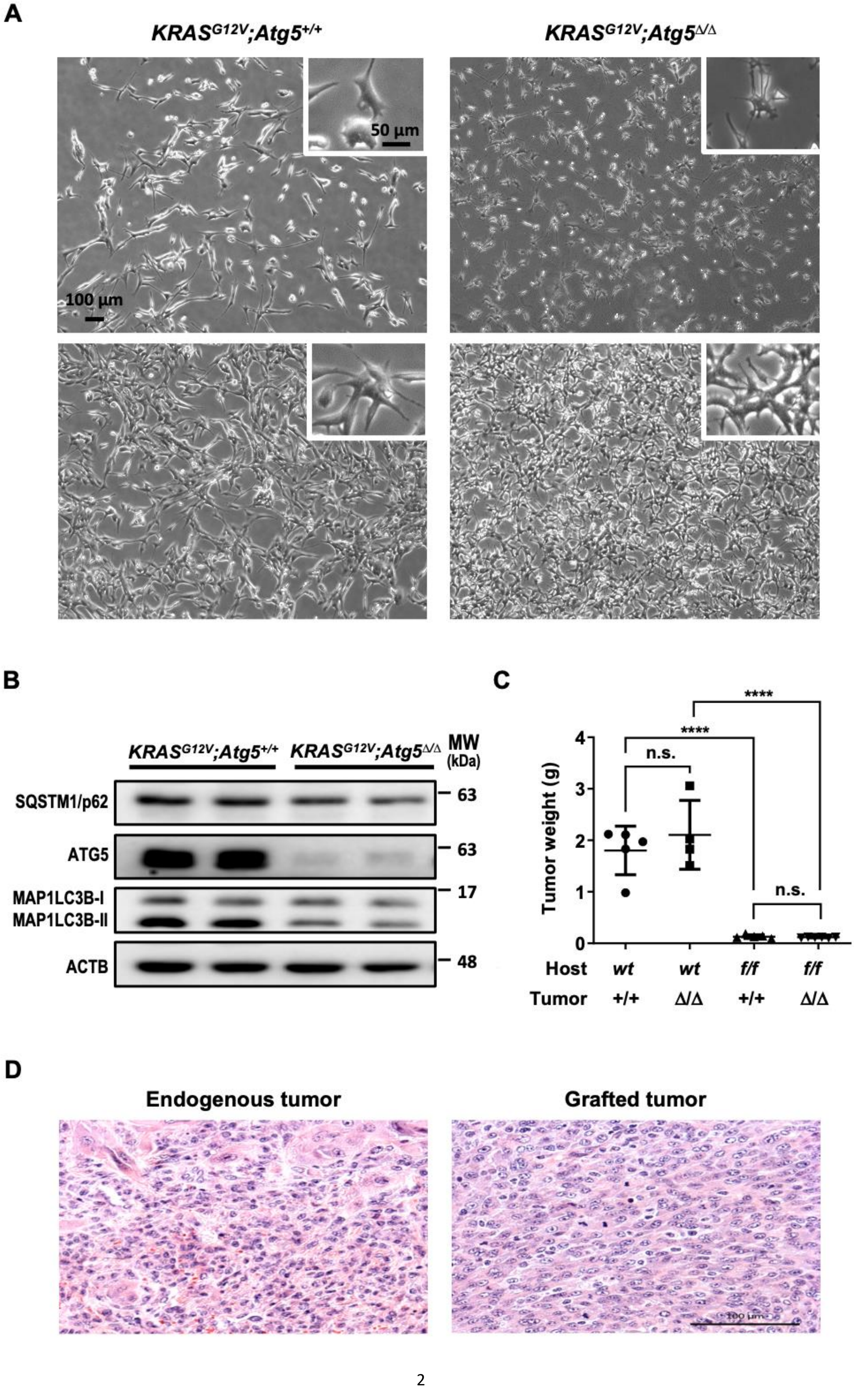
A
KRASG12V;Atg5+/+
KRASG12V;Atg5Δ/Δ
50 µm
100 µm
B
KRASG12V;Atg5+/+
KRASG12V;Atg5Δ/Δ
MW (kDa)
SQSTM1/p62
ATG5
MAP1LC3B-I
MAP1LC3B-II
ACTB
63
63
17
48
C
Tumor weight (g)
n.s.
****
****
n.s.
Host wt wt f/f f/f
Tumor +/+ Δ/Δ +/+ Δ/Δ
D
Endogenous tumor
Grafted tumor

**Supplemental Fig. S1. Characterization or primary salivary carcinoma cells isolated from different hosts.**

Primary tumor cells were isolated from tamoxifen-induced malignant salivary tumors. (**A**) Cell morphology of *KRAS*$^{G12V}$*;Atg5*$^{+/+}$ and *KRAS*$^{G12V}$*;Atg5*$^{\Delta/\Delta}$ malignant salivary tumor cells grown at low (top panels) and high (bottom panels) confluency. Enlarged images are also shown (inserts). (**B**) Representative western blots, in which total lysates from *KRAS*$^{G12V}$*;Atg5*$^{+/+}$ and *KRAS*$^{G12V}$*;Atg5*$^{\Delta/\Delta}$ malignant salivary tumor cells were stained with indicated antibodies. Two each of independent primary salivary tumor cells were isolated from tumor-bearing ATG5-wild type and ATG5-knockout mice, respectively, are shown. (**C**) Malignant salivary tumor cells *KRAS*$^{G12V}$*;Atg5*$^{+/+}$ (+/+) and *KRAS*$^{G12V}$*;Atg5*$^{\Delta/\Delta}$ (Δ/Δ) were orthotopically implanted into SMG of wild type (*wt*) and *Atg5*$^{flox/flox}$ (*f/f)* host mice. Tumor weights were measured on Day 25 post-implantation. $n \geq 4$. n.s., not significant; ****: $p < 0.0001$; Student's *t*-test, 2-tailed, unpaired. (**D**) Comparison between H&E stain of tumor sections of endogenous tamoxifen-induced salivary tumors and orthotopically implanted tumors. Scale bar; 100 μm.

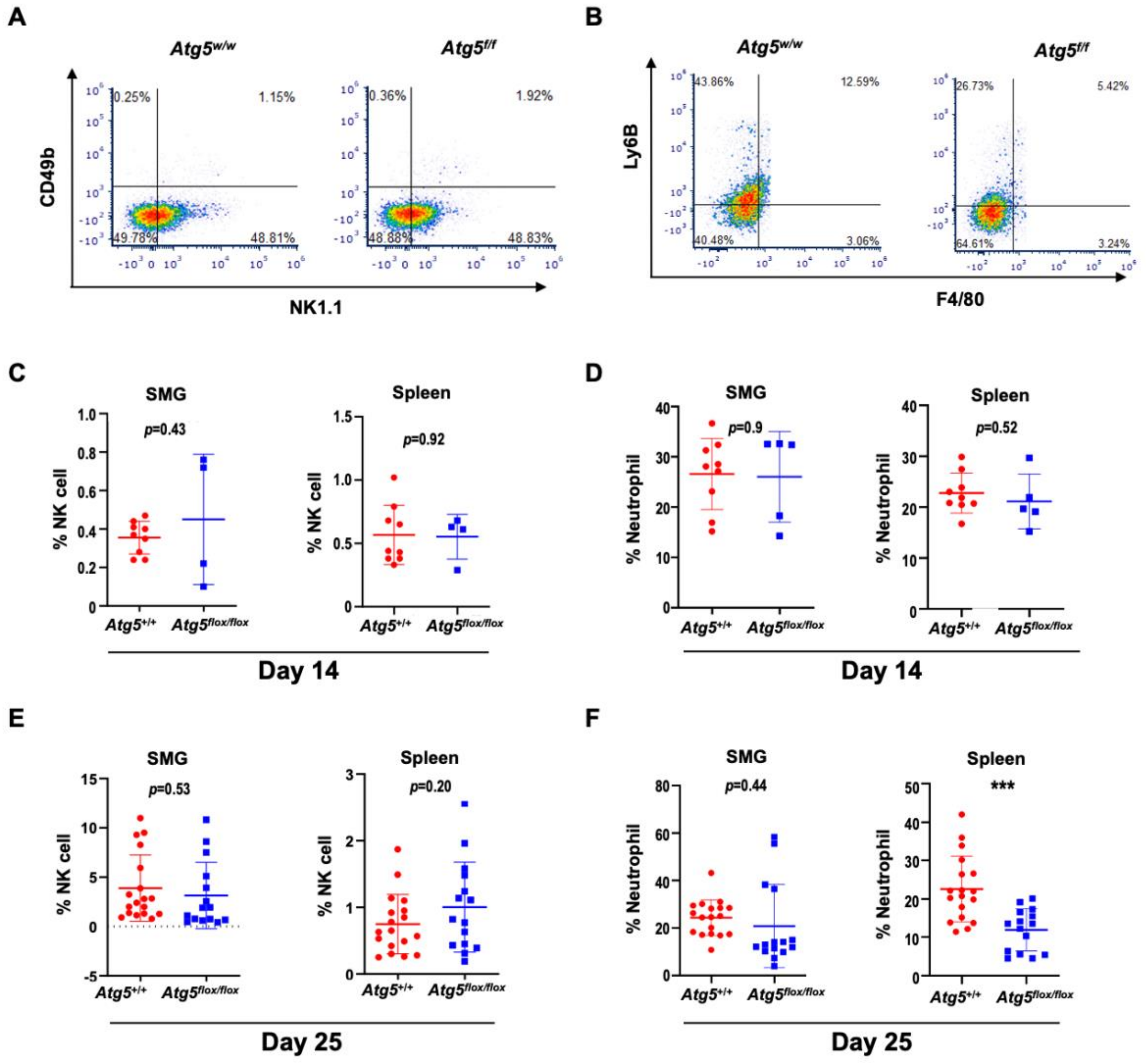


**Supplemental Fig. S2. Natural killer cells and neutrophils population in SMG TME.**

(**A**, **B**) Representative flow cytometry showing CD11b$^+$CD49b$^+$NK1.1$^+$ NK cells (**A**), and CD11b$^+$F4/80$^-$Ly6B$^+$ neutrophils (**B**) isolated from live splenocytes of Day 25 tumor-bearing *Atg5*$^{+/+}$ and *Atg5*$^{flox/flox}$ mice. (**C, E**) Flow cytometry analyses showing the percentage of NK cells (CD11b$^+$CD49b$^+$NK1.1$^+$) in alive SMG cells and splenocytes from Day 14 (**C**) and Day 25 (**E**) tumor-bearing *Atg5*$^{+/+}$ and *Atg5*$^{flox/flox}$ mice. (**D, F**) Flow cytometry analyses showing the percentage of neutrophils (CD11b$^+$F4/80$^-$Ly6B$^+$) in alive SMG cells and splenocytes from Day 14 (**D**) and Day 25 (**F**) tumor-bearing *Atg5*$^{+/+}$ and *Atg5*$^{flox/flox}$ mice. Day 14 tumor-bearing mice: *Atg5*$^{+/+}$: n = 9; *Atg5*$^{flox/flox}$: n = 5. Day 25 tumor-bearing mice: *Atg5*$^{+/+}$: n = 18; *Atg5*$^{flox/flox}$: n = 15. Data are shown as mean ± SD; ***: $p < 0.001$; Student's *t*-test, 2-tailed, unpaired.

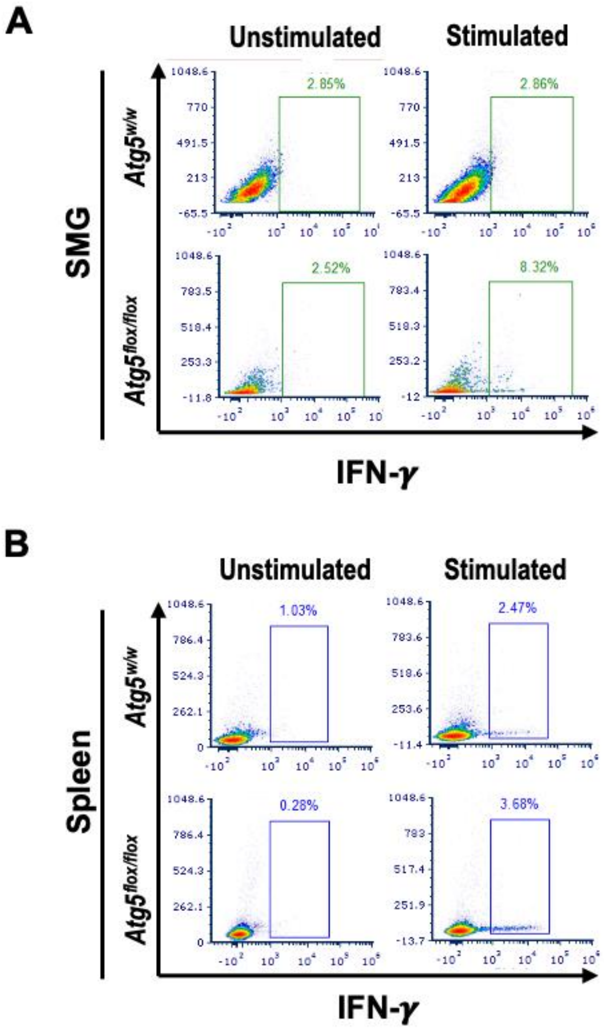


**Supplemental Fig. S3. Flow cytometry analysis of IFN-γ-producing cells in SMGs and spleens of tumor-bearing mice.**

Representative flow cytometry showing IFN-γ-producing cells in single cell suspensions of the alive SMG cells (**A**) and splenocytes (**B**) of tumor-bearing *Atg5*$^{+/+}$ and *Atg5*$^{flox/flox}$ mice following PMA/ionomycin/Golgiplug stimulation for 4h (Stimulated) or unstimulated control (Unstimulated).

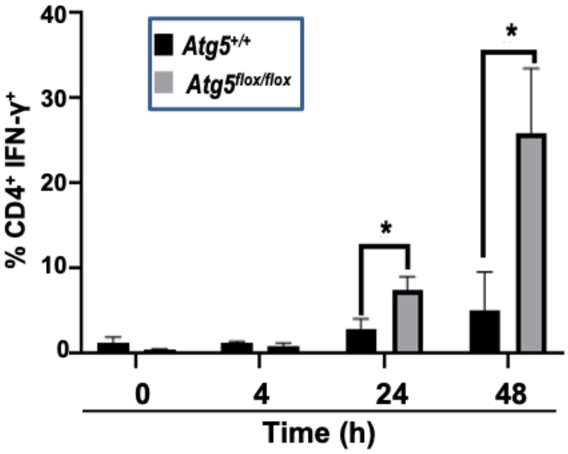


**Supplemental Fig. S4. Glutamine promotes IFN-γ$^{+}$ production by CD4$^{+}$ T cells from *Atg5$^{flox/flox}$* mice.**

CD4$^{+}$ T cells were isolated from splenocytes of naïve *Atg5$^{+/+}$* and *Atg5$^{flox/flox}$* mice and cultured in glutamine-free Treg polarization medium for 3 days. Afterwards, cells were cultured in fresh Treg polarization medium supplemented with glutamine (2 mM) for the indicated time periods (0, 4, 24 and 48 h), and then stimulated with PMA/ionomycin/Golgiplug for 4 h. Percentages of IFN-γ-positive CD4$^{+}$ cells were determined by flow cytometry analyses (n = 3). Data are presented in bar graph shown as Mean ± SEM. *p* value was calculated by *t* test (unpaired, two tailed). *: $p < 0.05$.

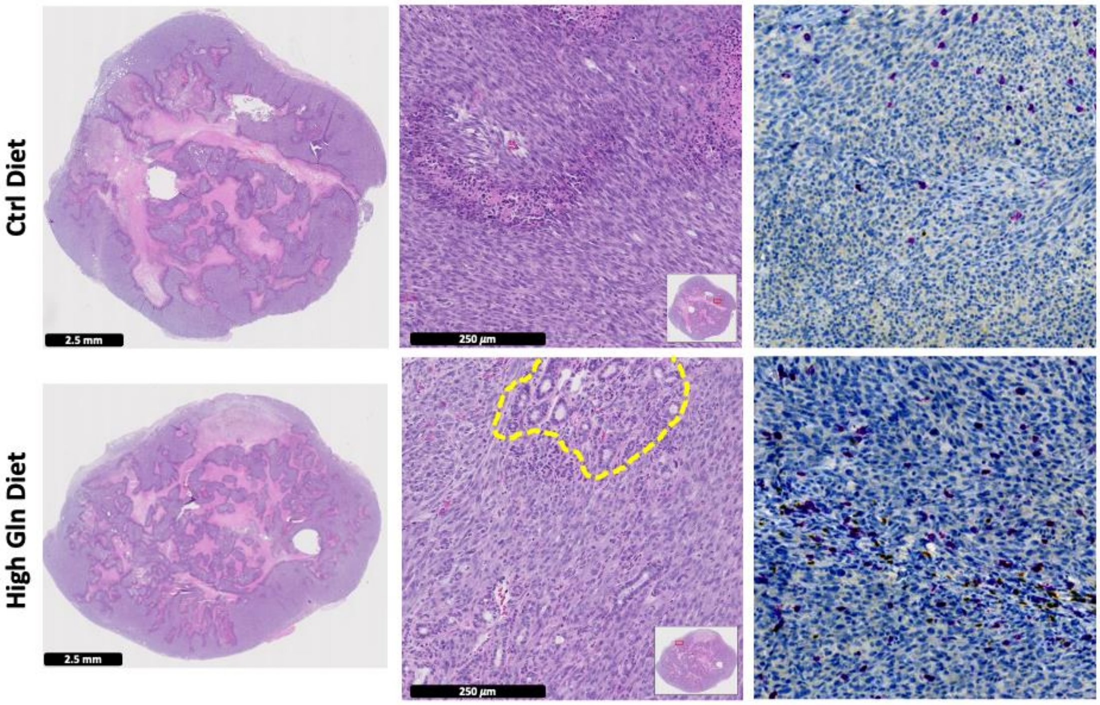


**Supplemental Fig. S5. High glutamine diet reduces MST tumor burden with increased CD8$^+$ T cell infiltration.**

Gross view (*left*) and enlarged view (*middle*) of H&E-stained tumor sections from two mouse cohorts, each fed with either a control (Ctrl) diet or high glutamine diet, respectively. Areas of residual salivary gland, *enclosed by a yellow dashed line,* can be seen in tumor from mice on a high glutamine diet. IHC staining with an anti-CD8 antibody shows increased tumor-infiltrating CD8 T cells (*purple in color*; *right*) in tumors from *Atg5*$^{+/+}$ mice fed with high-glutamine diet.

### Supplementary Table 1: Primer sets for RT-PCR analyses.

| Gene | Forward primer | Reverse primer |
|---|---|---|
| *Cdkn1a* | 5'- AGG AGC AAA GTG TGC CGT TG-3' | 5'- CGA AGT CAA AGT TCC ACC GTT C-3' |
| *Foxp3* | 5'- ACC ATT GGT TTA CTC GCA TGT -3' | 5'- TCC ACT CGC ACA AAG CAC TT -3' |
| *Gapdh* | 5'- CCC CTT CAT TGA CCT CAA CTA -3' | 5'- CTC CTG GAA GAT GGT GAT GG-3' |
| *Ifng* | 5'- CTT GAA CCC TGT CGT ATG CTG G -3' | 5'- TTG GTG CAG GAA TCA GTC CAG G -3' |
| *Il1a* | 5'- TGT TGC TGA AGG AGT TGC CAG -3' | 5'- CCC GAC TTT GTT CTT TGG TGG -3' |
| *Il1b* | 5'- TGG ACC TTC CAG GAT GAG GAC A -3' | 5'- GTT CAT CTC GGA GCC TGT AGT G-3' |
| *Il6* | 5'- TAC CAC TTC ACA AGT CGG AGG C-3' | 5'- CTG CAA GTG CAT CAT CGT TGT TC-3' |
| *Tnf* | 5'- GGT GCC TAT GTC TCA GCC TCT T -3' | 5'- GCC ATA GAA CTG ATG AGA GGG AG -3' |
| *Rps18* | 5'- TTC CAG CAC ATT TTG CGA GTA -3' | 5'- CAC GCC CTT AAT GGC AGT GAT -3' |
